# Highly Iterated Palindrome 1 (HIP1) sequence improves *Synechococcus* sp. PCC 7002 transformation efficiencies in a homology- and methylation-dependent manner

**DOI:** 10.1101/2024.09.10.611283

**Authors:** Cody Kamoku, David R. Nielsen

## Abstract

The ability to precisely control cyanobacterial metabolism first requires the ability to efficiently deliver engineered DNA constructs. Here, we investigate how natural transformation efficiencies in *Synechococcus* sp. PCC 7002 can be greatly improved by leveraging the native and abundant cyanobacterial Highly Iterated Palindrome 1 (HIP1) sequence. While including at least one homologous HIP1 site within the homology arms of an integrating plasmid increased integration efficiency by up to 7-fold, methylation of those sites by HIP1 methyltransferase (encoded by *slr0214* from *Synechococcus* sp. PCC 6803) boosted this to greater than a 100-fold improvement overall. Non-homologous HIP1 sites also improved transformation efficiencies of both integrating and replicating episomal plasmids (by up to 60- and 9-fold, respectively), but only when methylated. The collective data further reveal that HIP1 does not function as part of a native restriction enzyme system in PCC 7002, but rather may improve transformation efficiency via two complementary mechanisms: i) altering DNA binding/uptake/processing prior to homologous recombination, and ii) increasing the efficiency of homologous recombination in a manner reminiscent of a crossover hotspot instigator (Chi) site. Future studies are needed, however, to more clearly elucidate the specific role of HIP1 during natural transformation of cyanobacteria.

## 1. Introduction

Cyanobacteria use light energy to fix CO_2_ into biomass and thus show promise with respect to carbon capture and the sustainable bioproduction of chemicals^1–3^. Effectively doing so, however, will ultimately require high throughput strain engineering tools, such as CRISPRi/a, promoter, and/or gene libraries; the application of which currently remains bottlenecked by inadequate transformation efficiencies^4,5^. Since many commonly studied strains are naturally competent, natural transformation remains the most widely used mechanism for delivering DNA into cyanobacteria^6^. Current natural transformation methods are limited, however, by several inherent challenges. This includes the presence of endogenous nucleases, including restriction enzymes^7^ and/or CRISPR-Cas systems^8^ which natively function to defend against foreign DNA. Thus, as expected, removing native restriction sites from delivered DNA is one approach towards improving transformation efficiencies; with one study achieving a 60-fold improvement in PCC 7002 upon removing an AquI restriction site^9^. Meanwhile, other methods reported to improve natural transformation efficiencies in cyanobacteria include: *i)* deletion of the ssDNA exonucleases RecJ, which resulted in a ~100-fold enhancement in *Synechocystis* sp. PCC 6803 (PCC 6803)^10^; and, *ii)* addition of EDTA to inhibit extracellular nucleases, which resulted in an 8-fold increase in *Synechococcus elongatus* sp. PCC 7942 (PCC 7942)^11^.

Highly Iterated Palindrome 1 (HIP1) is an eight base pair sequence (consensus: GCGATCGC) that is highly abundant in virtually all cyanobacterial genomes, occurring on average every ~1000 bp in PCC 7002^12^. Many cyanobacteria with genomic HIP1 sites also contain predicted orphan methyltransferases that methylate the sequence^13^. One of these methylates a cysteine within HIP1 (G^m5^CGATCGC; henceforth referred to as HIP1 methyltransferase) and another an adenine in the GATC subsequence (G^m6^ATC; same target as *E. coli dam* methyltransferase)^13^, with both activities having been experimentally validated in PCC 6803^14,15^. In both PCC 6803 and *Anabaena* sp. PCC 7120, G^m6^ATC methylation is essential for growth^14,16^. Meanwhile, in *E. coli* and other bacteria, G^m6^ATC methylation is necessary for DNA mismatch repair, while deletion of *dam* has been found to increase mutation rates in PCC 7942^17^. In PCC 6803, HIP1 methyltransferase is encoded by *slr0214*, of which PCC 7002 also encodes a homolog (SYNPCC7002_A0849; 96% coverage, 50.5% identity). In PCC6803, deletion of *slr0214* is extremely detrimental under high light, high CO_2_ conditions^18^.

While the exact function(s) of HIP1 currently remains unknown, it has been hypothesized to play a role in DNA recombination/repair and/or maintenance of chromosome structure^12,13,19^. On the other hand, it has also been proposed that the role of HIP1 methyltransferase is to bypass a native restriction enzyme system targeting the HIP1 sequence^15^. Evidence for this hypothesis comes from the observation that natural transformation efficiencies of integration vectors containing HIP1 sites were improved by 11- to 161-fold in PCC 6803 following their methylation with HIP1 methyltransferase (i.e., miniprepping them from an *E. coli* strain expressing *slr2014*)^15^. That said, Scharnagl et al. earlier reported that PCC 6803 does not possess restriction enzyme activity targeting unmethylated HIP1 sites^18^. Moreover, if this were indeed true, in the traditional paradigm of restriction modification systems, complete removal of the restriction site should also increase transformation efficiency whereas addition of an unmethylated site should reduce it. On the contrary, here we demonstrate how unmethylated HIP1 sites instead increase transformation efficiencies in PCC 7002, but only when they share homology with a native chromosomal HIP1 site. Methylation by HIP1 methyltransferase, meanwhile, significantly enhances the effect. Furthermore, as demonstrated for both chromosomal integration and episomal replicating vectors, non-homologous HIP1 sites also increase transformation efficiency, but only when methylated by HIP1 methyltransferase. Taken together, our collective findings suggest that HIP1 may enhance natural transformation in PCC 7002 via multiple, complementary roles. Ultimately, this study provides an easily implemented methodology for improving transformation efficiencies in PCC 7002 and likely other cyanobacteria, adoption of which will greatly enhance the overall design-build-test-learn (DBTL) cycle.

## Materials and methods

### Strain construction

All strains constructed and/or used in this study are summarized in Table 1. Plasmid pAC-CV (a gift from Prof. Deirdre Meldrum) served as the template for cloning *slr0214* from PCC 6803^15^. Following construction of pCMK83 and pKDsgRNA gidB-3, *E. coli* DHIP1 was constructed by inserting *slr0214* into the *gidB-atpI* intergenic locus of *E. coli* DH10β using the CRISPR-based “no-SCAR” method^20^. In this case, as in pAC-CV, *slr0214* was constitutively expressed via the promoter of the Tetracycline resistance gene. PCC 7002 Δ*addAB* was constructed using pCJ266 to replace the *addAB* (SYNPCC7002_F0007, SYNPCC7002_F0008) locus of PCC 7002 with a gentamicin resistance cassette via homologous recombination.

**Table 1.** Strains and plasmids constructed and/or used in this study.

| Strain | Relevant Characteristics | Source |
| --- | --- | --- |
| *E. coli* DH10β | $\Delta(mrr\text{-}hsdRMS\text{-}mcrBC)$ | Invitrogen |
| *E. coli* DHIP1 | DH10β *atpI::slr0214::gidB* | This study |
| *Synechococcus* sp. PCC 7002 | Wild type | UTEX |
| *Synechococcus* sp. PCC 7002 $\Delta addAB$ | $\Delta addAB$ ($\Delta SYN PCC 7002\_F0007$, $\Delta SYN PCC 7002\_F0008$) | This study |
| Plasmid | Relevant Characteristics | Source |
| pAC-Cv | pACYC184 (Cmr Tetr, p15A ori) expressing *slr0214* from PCC 6803 | Wang et al.<sup>15</sup> |
| pKDsgRNA *gidB*-3 | gRNA targeting *gidB-atpI* locus of *E. coli* | This study |
| pCMK83 | Integrates *slr0214* into *gidB-atpI* locus of *E. coli* | This study |
| pCJ266 | Integrates *Gm<sup>R</sup>* into *addAB* ($SYN PCC 7002\_F0007$, $SYN PCC 7002\_F0008$) | This study |
| pCMK84 | Integrates *Kan<sup>R</sup>* into NS2<sup>23</sup>, 250 bp homology, no HIP1 site | This study |
| pCMK85 | pCMK84 with a homologous HIP1 site included at the 3' end of right homology arm | This study |
| pCMK95 | pCMK84 with a non-homologous HIP1 site included before the 5' end of the left homology arm | This study |
| pCMK100 | pCMK84 with a non-homologous HIP1 site included in the plasmid backbone | This study |
| pCMK103 | Integrates *Kan<sup>R</sup>* into NS1, 500 bp homology, one homologous HIP1 site in left homology arm | This study |
| pCMK104 | Integrates *Kan<sup>R</sup>* into NS2, 500 bp homology, one homologous HIP1 site in right homology arm | This study |
| pCMK105 | Integrates *Kan<sup>R</sup>* into *glpK*, 570 bp homology, three homologous HIP1 sites in left homology arm | This study |
| pCMK139 | Shuttle vector (pSC101 ori, pCB2.4 ori, *Kan<sup>R</sup>*) that replicates in *E. coli* and PCC 7002 and contains one non-homologous HIP1 site | Kamoku et al.<sup>24</sup> |
| pCMK138 | pCMK139 with HIP1 site removed | This study |
| pCMK179 | pCMK84 with left homology arm extended to 790 bp | This study |
| pCMK180 | pCMK179 with a homologous HIP1 site included at the 3' end of left homology arm | This study |
| pCMK181 | pCMK179 with a homologous HIP1 site included at the 5' end of left homology arm | This study |
| pCMK182 | pCMK179 with homologous HIP1 sites included at the 5' end of left homology arm and 3' end of right homology arm | This study |
| pCMK195 | Integrates *Kan<sup>R</sup>* into NS2, 1250 bp homology, two homologous HIP1 sites in each of the right and left homology arms | This study |

### Plasmid construction

All plasmids constructed and/or used in this study are summarized in Table 1. Genomic DNA from PCC 7002 served as the template for generating plasmids containing homology arms for integration into different chromosomal loci of interest. Integration plasmids were comprised of the pBR322 origin of replication together with a *Kan^R^* cassette (for selection of transformants) flanked by homology arms (each 250-1250 bp) specific to the target integration site. The pCMK138 replicating plasmid was constructed by removing the HIP1 site from pCMK139 via Golden Gate assembly^21^. All primers were custom designed and synthesized by IDT (Supplemental Table 1). DNA fragments were PCR amplified using Q5 High Fidelity Polymerase (NEB). Integrating plasmids were constructed via Golden Gate assembly^21^ utilizing BsaI (NEB), PaqcI (NEB), or BpiI (Thermo Fisher Scientific). Alternatively, plasmids were constructed using Gibson Assembly^22^.

### Preparation of methylated and unmethylated plasmids

All plasmids were prepared from 25-50 mL overnight cultures grown at 37°C in LB with appropriate antibiotics and purified using the ZymoPURE II Plasmid Midiprep Kit (Zymo Research). Plasmids with methylated HIP1 sites were miniprepped from *E. coli* DHIP1 whereas those with unmethylated HIP1 sites were miniprepped from *E. coli* DH10β. Plasmids miniprepped from both DHIP1 and DH10β contain methylated Dam sites. Plasmids with both unmethylated HIP1 and Dam sites were miniprepped from *dam*-/*dcm*-*E. coli* (NEB). To obtain plasmids with only methylated Dam sites (and not HIP1), plasmids miniprepped from *dam*-/*dcm*-*E. coli* were treated in vitro with Dam Methyltransferase (NEB). To obtain plasmids with only methylated HIP1 sites (and not Dam), the plasmid of interest was co-transformed with pAC-Cv into *dam*-/*dcm*-*E. coli*. The plasmid of interest was then isolated from the miniprepped mixture via gel extraction. Plasmids with both methylated HIP1 and Dam sites were alternatively generated via in vitro treatment of the above gel extracted plasmids with Dam Methyltransferase. Miniprepped replicating plasmids were fractionated via gel electrophoresis after which individual bands enriched with multimers were extracted, as previously described^24^.

### Transformation of PCC 7002

Plasmids designed for chromosomal integration were transformed as follows. PCC 7002 was grown overnight at 37°C in 250 mL shake flasks containing 15-20 mL Media A+ supplemented with 30 μM ferric ammonium citrate^25^, while shaking at 220 RPM and with 150 μE light intensity under a 1% CO_2_ atmosphere. The next morning, by which time cell growth had reached an OD_730_ of 0.5-0.7, cells were removed by centrifugation at 3000 x *g*, and the pellet resuspended in fresh Media A+ to an OD_730_ of 0.6. Next, 100-1000 ng of plasmid DNA was added to 1 mL of resuspended cells in a 15 mL culture tube and then incubated for 24 h at 37°C while shaking at 220 RPM with 150 μE light intensity and atmospheric CO_2_. Different volumes of cells (100 – 900 μL; as appropriate, based on the expected transformation efficiency of the experiment) were then plated on Media A+ agar plates (0.75% w/v^26^) with 100 μg/mL kanamycin. Plates were incubated at 37°C with 150 μE light intensity and atmospheric CO_2_ for 6-7 days before the resulting colonies were then counted. Transformation of replicating plasmids was carried out in the same manner, except that cells were resuspended in fresh Media A+ to an OD_730_ of 1.0 and incubated for 3 hours before plating on Media A+ agar plates (0.75% w/v) with 75 μg/mL kanamycin. For experiments comparing integration vectors targeting the same genomic loci, transformation efficiency was determined by counting the resulting number of kanamycin-resistant colonies and dividing by both the volume of cells plated and the mass of DNA added. Results were then normalized by dividing each resulting transformation efficiency by the average transformation efficiency of the control (plasmid with no HIP1 site), which was included in each experiment. For experiments comparing integration vectors that target different genomic loci or experiments comparing the transformation of two different strains of PCC 7002, transformants were plated both with and without kanamycin. Transformation efficiency was then determined by dividing the number of kanamycin-resistant colonies by the total number of colonies obtained without antibiotic.

## Results

### Introducing and verifying HIP1 methyltransferase activity in *E. coli* DHIP1

To generate plasmids with methylated HIP1 sites, *E. coli* DHIP1 was first constructed by integrating and constitutively expressing *slr0214* from PCC 6803 from the *gidB-atpI* intergenic locus of *E. coli* DH10β. HIP1 methyltransferase activity was confirmed by restriction digest treatment with PvuI, which targets the CGATCG sequence internal to the HIP1 site. In this case, PvuI digestion is blocked due to cytosine methylation by HIP1 methyltransferase. As seen in Supplementary Figure 1, plasmid pCMK85 miniprepped from *E. coli* DHIP1 was protected from PvuI digestion whereas pCMK85 miniprepped from *E. coli* DH10β was digested.

### Methylation of HIP1 sites increases transformation efficiency across multiple genomic loci

To investigate if HIP1 methylation improves transformation of PCC 7002 and if those effects are generalizable across multiple chromosomal loci, three different integrating plasmids were constructed, each targeting a commonly used integration site in PCC 7002. Since HIP1 sites occur on average every ~1000 bp in PCC 7002 and homology arms designed for chromosomal integration are typically ~500 bp in length, there is high likelihood that one or more native HIP1 sites will be present within a given integration plasmid design. As seen in Figure 1A, this was indeed the case with respect to each of pCMK103, pCMK104 and pCMK105; plasmids designed for integration into the frequently used neutral site 1 (NS1), neutral site 2 (NS2) and *glpK* (encoding glycerol kinase, which is dispensable during photoautotrophic growth) integration sites, respectively^23,27^. In this case, pCMK103 contained one HIP1 site in the left homology arm, pCMK104 contained one HIP1 site in the right homology arm, and three HIP1 sites were present in pCMK105, all in the left homology arm (Figure 1A). As shown in Figure 1B, HIP1 methylation enhanced transformation efficiencies across all three sites, representing up to a 7.5-fold improvement overall. Meanwhile, despite including three homologous HIP1 sites in pCMK105, no greater effect on transformation efficiency was realized, suggesting that the presence of a single methylated HIP1 site was sufficient.

**Figure 1.**
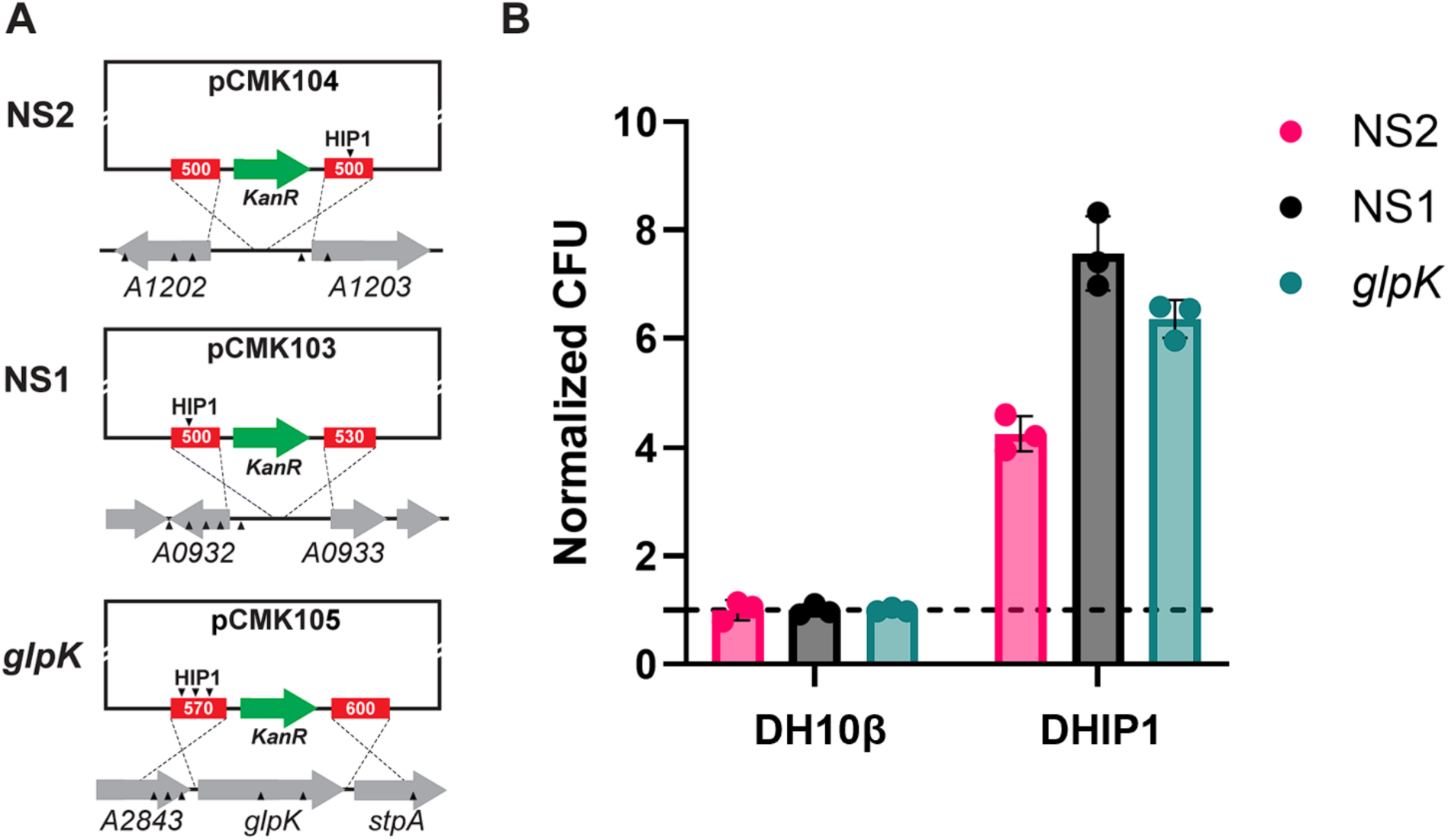
Methylation of HIP1 sites in homology arms improves natural transformation efficiencies across three distinct chromosomal loci. A: Schematic illustrating the number and positioning of native HIP1 sites located in the homology arms designed for each locus. All plasmids integrate a Kan^R^ cassette into their corresponding chromosomal locus. Solid black triangles represent native and homologous HIP1 sites. B: Comparison of transformation efficiencies obtained when delivering HIP1 methylated or unmethylated pCMK103 (NS1), pCMK104 (NS2), and pCMK105 (glpK). Transformation efficiency measured as the number of kanamycin-resistant colonies divided by the total number of colony forming units (CFUs). ‘Relative CFUs’ determined by dividing the results for each locus by those of the HIP1 unmethylated condition (i.e., dam+ HIP1-). Error bars represent standard deviation from triplicate experiments.

### Including and methylating a homologous HIP1 site enhances transformation efficiency

With methylation by HIP1 methyltransferase resulting in enhanced transformation efficiencies, the results in Figure 1 so far do not refute the possibility that HIP1 indeed functions by blocking a native restriction enzyme system. To more thoroughly explore this potential mechanism, we next sought to examine how the presence of a HIP1 site, regardless of its methylation state, affects transformation efficiency in PCC 7002. To do so, another integration vector targeting NS2 (pCMK84) was constructed wherein the homology arms were truncated (each just 250 bp) to omit the naturally occurring HIP1 site originally located in the right homology arm of pCMK104 (Figure 2A). The right homology arm of pCMK84 was then extended by 8 bp to reintroduce the same native HIP1 site, resulting in pCMK85 (Figure 2A). Here, since total homology remained 97% conserved between pCMK84 and pCMK85, it was reasoned that any differences would be attributable to the HIP1 site itself and not an artifact caused by using slightly longer homology arm^23^. As shown in Figure 2B, when unmethylated, inclusion of a HIP1 site in the homology region of pCMK85 resulted in a 7-fold increase in transformation efficiency relative to pCMK84; a finding that stands in contrast to the hypothesis that HIP1 functions as a restriction site. Lastly, and most significantly, methylation of the homologous HIP1 site in pCMK85 further boosted transformation efficiency by an additional ~12-fold, collectively representing an 83-fold improvement over pCMK84 (Figure 2B).

**Figure 2.**
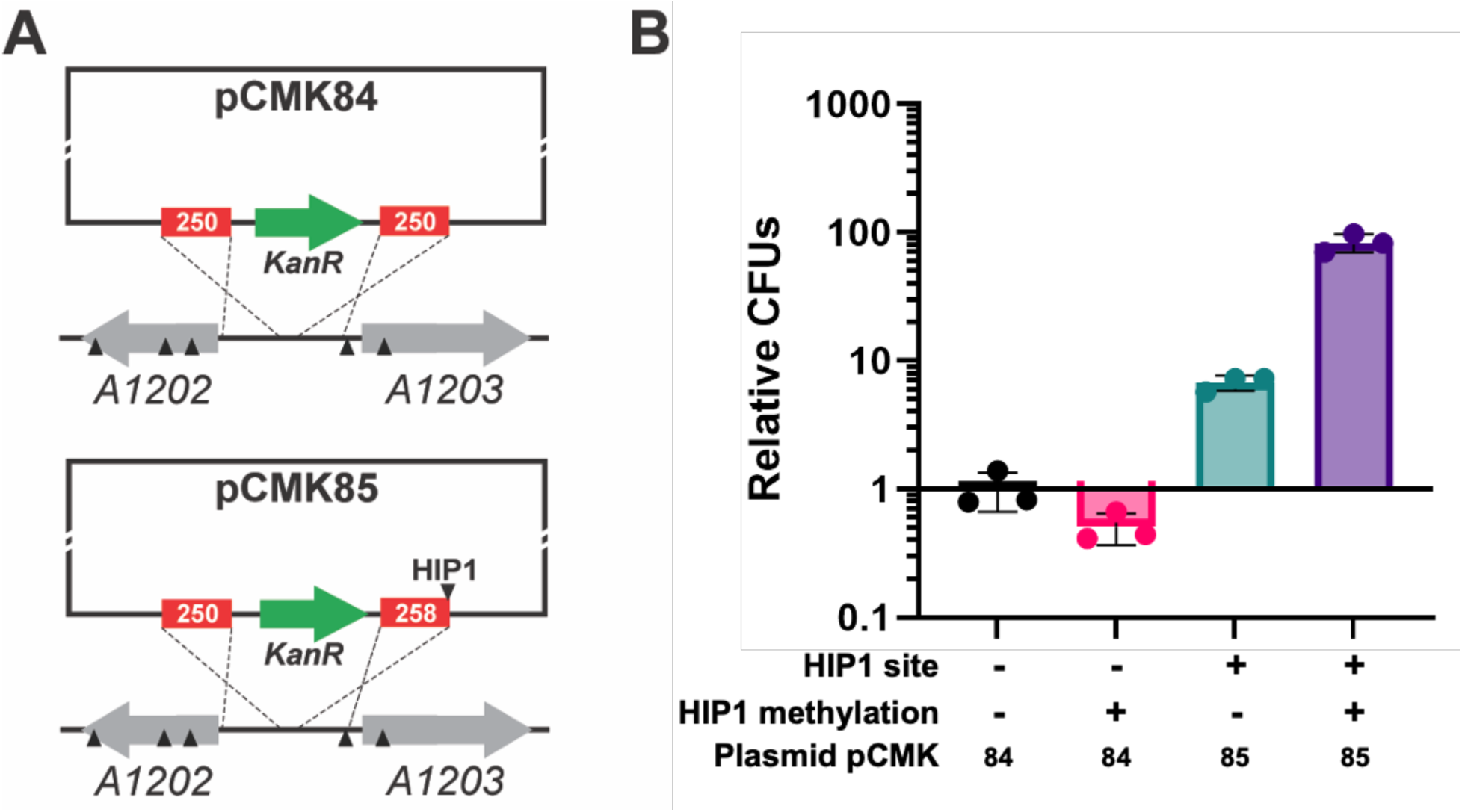
Inclusion and methylation of a homologous HIP1 site enhances transformation efficiency in PCC 7002. A: Schematic illustrating the number and positioning of native HIP1 sites (if any) located in the homology arms of pCMK84 and pCMK85. Both plasmids integrate a Kan^R^ cassette into the NS2 chromosomal locus. Solid black triangles represent native and homologous HIP1 sites. B: Effect of including and methylating a HIP1 site on the relative number of colony forming units (CFUs) obtained For all plasmids, CFUs were first normalized by dividing by the volume of cells plated and mass of DNA transformed. ‘Relative CFUs’ were determined by comparing the normalized results to those obtained for pCMK84. Error bars represent standard deviation from triplicate experiments.

As discussed above, HIP1 sites contain two potential methylation sites: *i)* the first cytosine, which is the target of HIP1 methyltransferase (G^5m^CGATCGC), and *ii)* the adenine within the GATC subsequence, which is the target of Dam methyltransferase (GCG^6m^ATCGC)^13^. This is notable since both *E. coli* DHIP1 and DH10β encode *dam*, which could confound our understanding of the effects of methylation status on transformation efficiency. Therefore, we next assayed both the individual and combined effects of HIP1 and Dam methyltransferase on transformation efficiency, in this case using pCMK85. However, as shown in Supplemental Figure 2, Dam methylation provided little to no benefit towards transformation efficiency, either alone or in combination with HIP1 methylation. As such, all subsequent experiments examining the effects of HIP1 methylation were performed using plasmids miniprepped from either DH10β (*dam*+ HIP1-) or DHIP1 (*dam*+ HIP1+).

### Additional homologous HIP1 sites and longer homology arms further improve transformation efficiency

Although the outcomes of Figure 1 demonstrate how including a methylated HIP1 site within either homology arm enhances transformation efficiency, since that experiment spanned multiple chromosomal loci, no definitive conclusions could be drawn with respect to the potential influence of positional bias and/or optimal positioning of homologous HIP1 sites. Furthermore, as those HIP1 sites were internal to their respective homology arms, simply removing them would disrupt homology, thereby obfuscating the specific effects of HIP1. To clarify these points, the left homology arm of pCMK84 was next extended to 790 bp, stopping just short of a native HIP1 site (pCMK179). Homologous HIP1 sites were then introduced into the left (pCMK180), right (pCMK181), or both (pCMK182) homology arms by simply extending them 8 bp in the appropriate direction(s) (Figure 3A). As shown in Figure 3B, including and methylating a single homologous HIP1 site in either homology arm resulted in nearly a 100-fold increase in transformation efficiency (relative to no HIP1 site; pCMK179). Doing so in both homology arms resulted in a 130-fold increase in transformation efficiency (relative to no HIP1 site; pCMK179). Although only a 1.3-fold increase in transformation efficiency compared to a single homologous HIP1 site in either arm, this increase was nevertheless statistically significant (p = 0.0016 for pCMK182 vs. pCMK180 and p < 0.0001 for pCMK182 vs. pCMK181; unpaired t-test with Welch’s correction).

**Figure 3.**
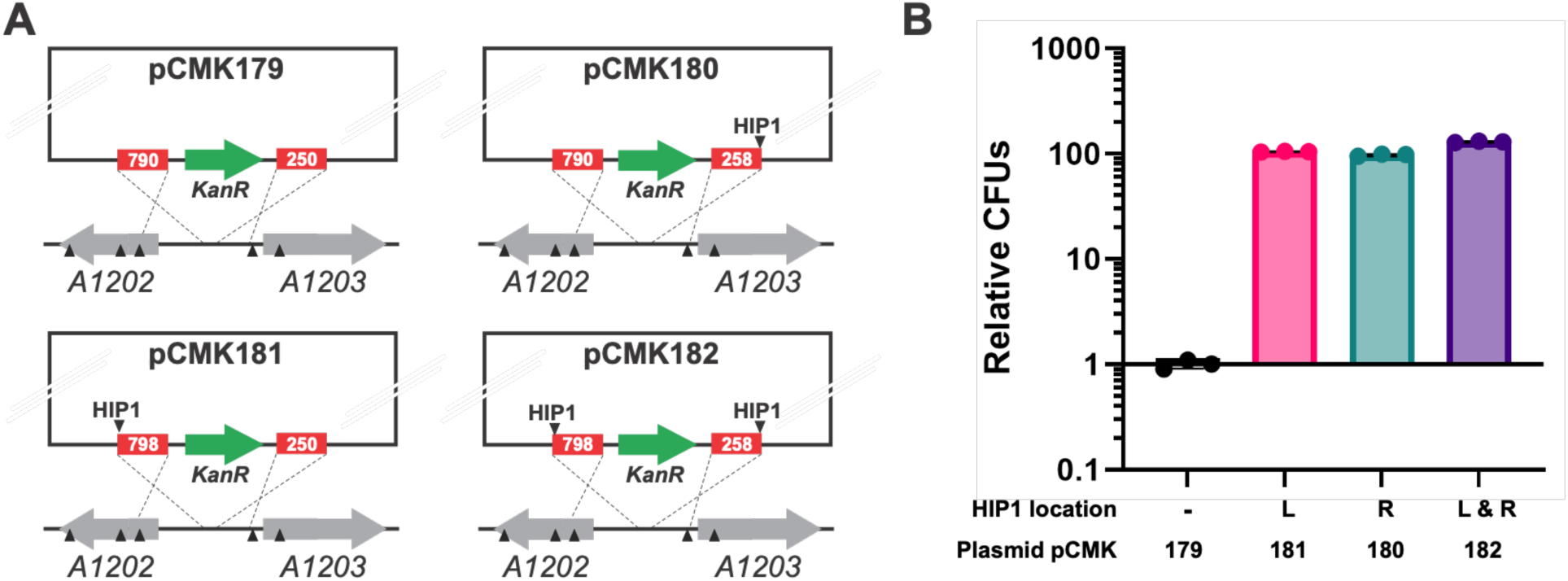
Positioning and total number of homologous HIP1 sites does not significantly affect transformation efficiency in PCC 7002. A: Schematic illustrating the number and relative positioning of homologous HIP1 sites located in the homology arms of pCMK179, pCMK180, pCMK181, and pCMK182. All plasmids integrate a Kan^R^ cassette into the NS2 chromosomal locus and were miniprepped from E. coli DHIP1. Solid black triangles represent native and homologous HIP1 sites. B: Effect of including and methylating homologous HIP1 sites in different positions within each homology arm on the relative number of colony forming units (CFUs) obtained. For all plasmids, CFUs were first normalized by dividing by the volume of cells plated and mass of DNA transformed. ‘Relative CFUs’ were determined by comparing the normalized results to those obtained for pCMK179. Error bars represent standard deviation from triplicate experiments.

Lastly, as previously reported by Ruffing et al., transformation efficiencies in PCC 7002 are also dramatically improved by increasing the length of both homology arms up to 1250 bp^23^. Accordingly, pCMK195 was next constructed with 1250 bp homology arms to explore the effects of combining this approach with the inclusion and methylation of HIP1 sites. Like pCMK85 and pCMK104, whose respective homology arms were ~250 bp and 500 bp in length, pCMK195 also integrates into the NS2 neutral site. Differently, however, pCMK195 contains four homologous HIP1 sites (two in each homology arm) versus just one for both pCMK85 and pCMK104 (Figure 4A). As shown in Figure 4B, transformation efficiency was indeed found to be a function of homology arm length and, by increasing them from 500 bp to 1250 bp, a ~100-fold enhancement was realized; here reaching a maximum of 2.4 x 10^7^ CFUs/μg DNA (or 5.11 x 10^-2^ Kan^R^ CFUs/total viable CFUs).

**Figure 4.**
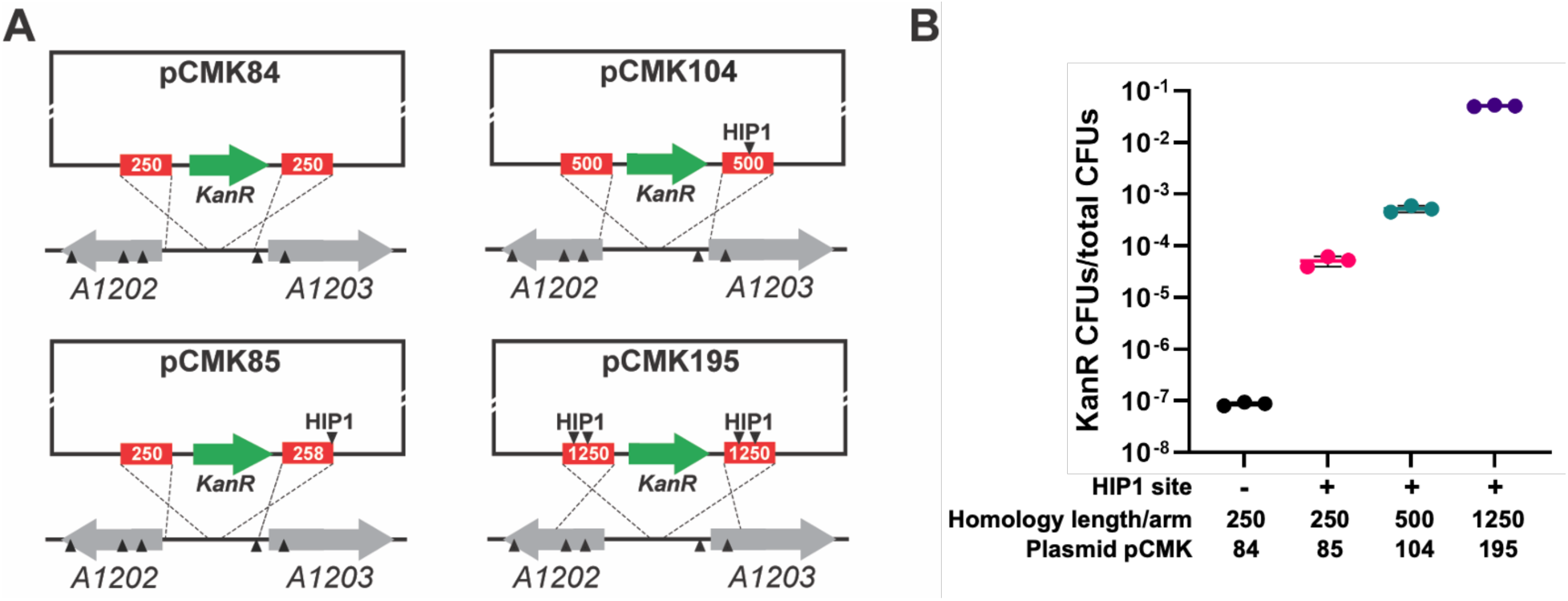
Longer homology arms support increased transformation efficiencies in PCC 7002. A: Schematic illustrating the number and relative positioning of homologous HIP1 sites located in the homology arms of pCMK84, pCMK85, pCMK104, and pCMK195, along with the length and relative positioning of homology arms. All plasmids integrate a Kan^R^ cassette into the NS2 chromosomal locus. Solid black triangles represent native and homologous HIP1 sites. B: Effect of homology arm length on number of Kan^R^ colony forming units (CFUs) obtained, as a proportion of all total viable CFUs. For all plasmids, Kan^R^ CFUs were counted then divided by the total number of CFUs when plated on no antibiotic plates. Error bars represent standard deviation from triplicate experiments.

### Non-homologous HIP1 sites enhance transformation efficiency when methylated

Thus far, the effects of including and methylating HIP1 sites have exclusively focused on their positioning within one or both homology arms of the integrating plasmid. To probe the potential importance of homology, a series of integrating plasmids was next constructed wherein non-homologous HIP1 sites were synthetically added external to the integration cassette (Figure 5A). This included pCMK100, with a HIP1 site added directly adjacent to the left homology arm, and pCMK95, with it added at a distal position. The results, compared in Figure 5B, reveal several important findings. First, unlike for homologous HIP1 sites (Figure 2B), unmethylated non-homologous HIP1 sites provided little to no benefit to transformation efficiency (note: although statistically significant, the difference between unmethylated pCMK95 and pCMK84 was only 3.4-fold; unpaired t-test with Welch’s correction, p=0.0006). When methylated, however, non-homologous HIP1 sites supported up to 60-fold greater transformation efficiencies relative to a control without a HIP1 site (pCMK84).

**Figure 5.**
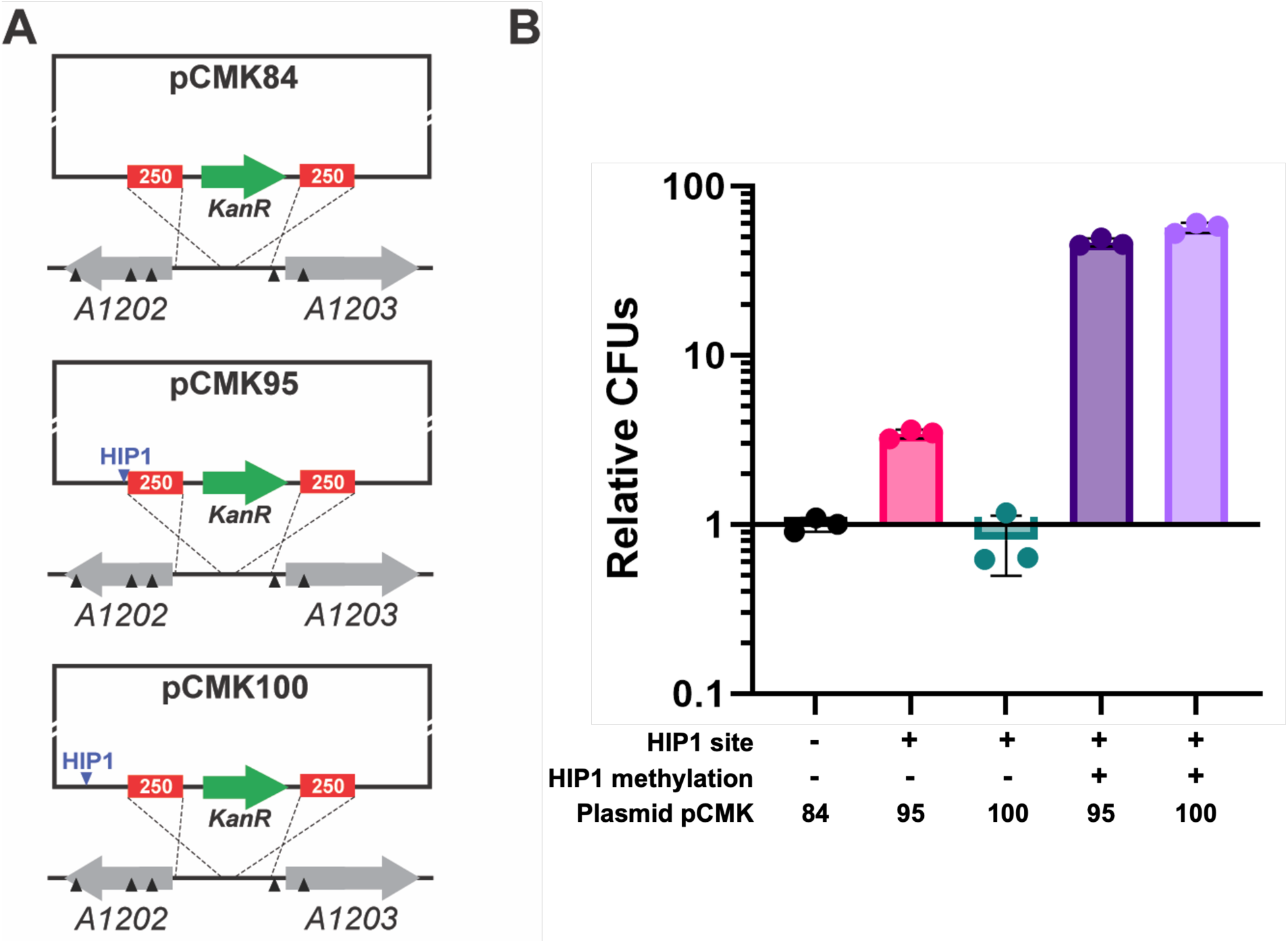
Non-homologous HIP1 sites increase transformation efficiency in PCC 7002 when methylated. A: Schematic illustrating the number and positioning of non-homologous HIP1 sites in pCMK95 and pCMK100. Both plasmids integrate a Kan^R^ cassette into the NS2 chromosomal locus. Solid black triangles represent native and homologous HIP1 sites. Solid blue triangles represent non-homologous HIP1 sites. B: Effect of including and methylating non-homologous HIP1 sites on the relative number of colony forming units (CFUs) obtained. For all plasmids, CFUs were first normalized by dividing by the volume of cells plated and mass of DNA transformed. ‘Relative CFUs’ were determined by comparing the normalized results to those obtained for pCMK84. Error bars represent standard deviation from triplicate experiments.

The influence of non-homologous HIP1 sites and their methylation on transformation efficiency was next further explored via the use of replicating episomal plasmids. Unlike the integrating plasmids investigated thus far, replicating plasmids share no homology with PCC 7002, thereby allowing the effects of non-homologous HIP1 sites to be studied independently from any putative role that HIP1 might play with respect to homologous recombination. For this purpose, we utilized pCMK139; a shuttle vector recently engineered by our group that carries both a pSC101 origin of replication and the pCB2.4 origin of replication from PCC 6803 (Figure 6A)^24^. pCMK139 encodes a single HIP1 site and can be delivered to PCC 7002 via natural transformation in the form of plasmid multimers. For comparison, pCMK138 was engineered here by removing the HIP1 site. Dimeric fractions of pCMK138 and pCMK139 were isolated by gel extraction of samples miniprepped from *E. coli* DH10β or DHIP1 and then transformed into PCC 7002. In this case, as shown in Figure 6B, the non-homologous HIP1 site in pCMK138 benefitted transformation efficiency only when methylated; a result consistent with the outcomes of Figure 5. Specifically, replicating plasmid transformation efficiency was improved up to ~9-fold by including and methylating a single non-homologous HIP1 site.

**Figure 6.**
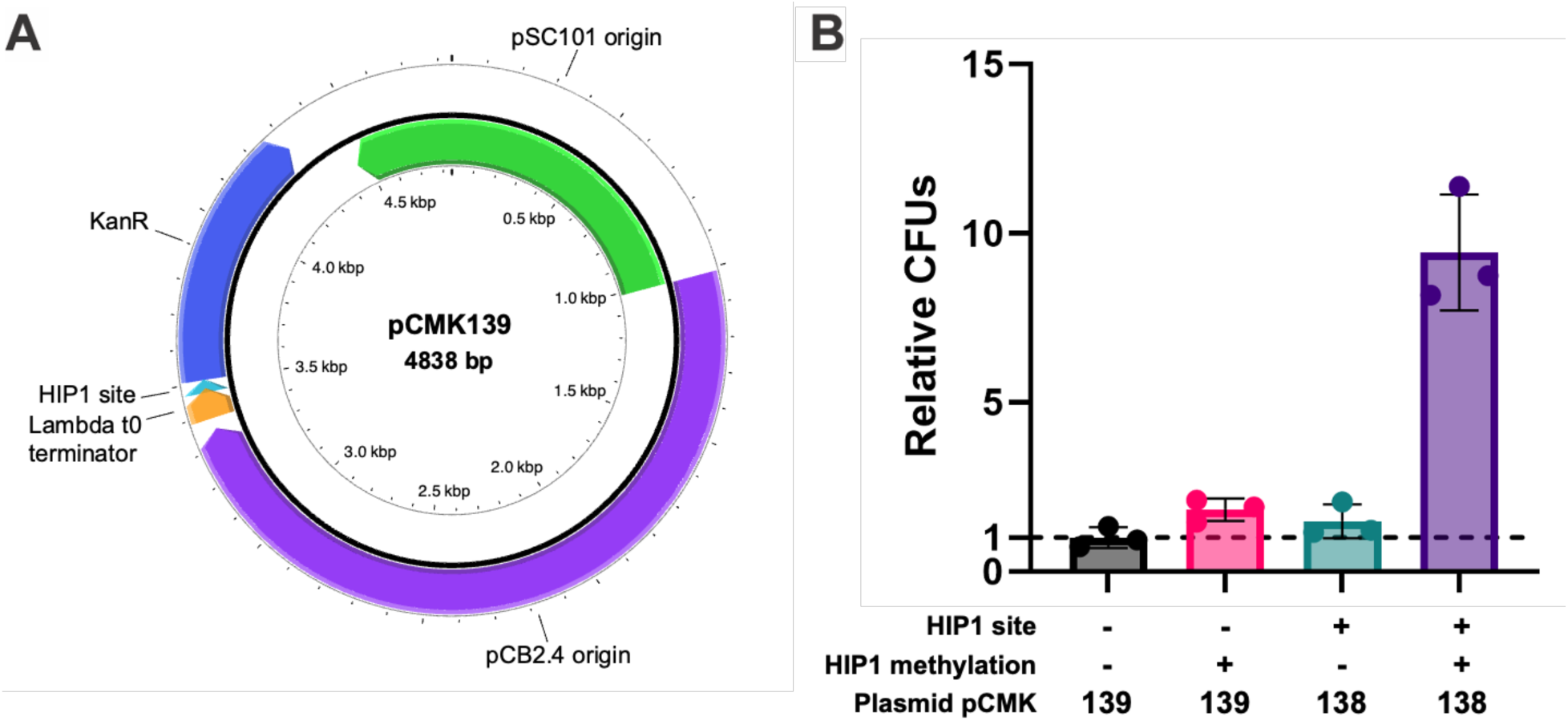
Including and methylating a non-homologous HIP1 site enhanced the transformation efficiency of replicating plasmids. A: Plasmid map of the pCMK138 replicating vector used in this experiment. The HIP1 site was removed to create pCMK138 (not shown). Plasmid map was generated using PlasMapper 3.0^28^. B: Effect of including and methylating a non-homologous HIP1 site on the relative number of colony forming units (CFUs) obtained. Both plasmids were transformed in their multimeric form. For all plasmids, CFUs were first normalized by dividing by the volume of cells plated and mass of DNA transformed. ‘Relative CFUs’ were determined by comparing the normalized results to those obtained for pCMK139 miniprepped from DH10β. Error bars represent standard deviation from triplicate experiments.

### Investigating alternative mechanisms of HIP1 function

If not a restriction site, one alternative explanation for the function of HIP1 is that of a cyanobacterial Chi (crossover hotspot instigator) site analogue. Chi sites serve as recombination hotspots in *E. coli* and other bacteria, and HIP1 was previously proposed as a recombination hotspot in PCC 7002^19^. In many bacteria, following a double strand break, the nuclease/helicase complexes RecBCD or AddAB are responsible for unwinding DNA. This occurs until they encounter a Chi site, at which point the complex cuts one strand to produce ssDNA, after which RecA is recruited to promote recombination and repair the damaged DNA^29,30^. While PCC 7002 does encode a homolog to RecA, homologs of RecC and RecD are missing^31^. PCC 7002 does, however, encode homologs of *Bacillus subtilis addA* and *addB* within an operon on the native pAQ6 plasmid (*addA*: SYNPCC7002_F0008 (73% coverage, 25.8% identity), GenBank accession ACB00939.1; *addB*: SYNPCC7002_F0007 (16% coverage, 21.4% identity), GenBank accession ACB00938.1). As a cursory test of HIP1’s alternative function as a Chi site, these *addAB* homologues were first knocked out in PCC 7002, resulting in PCC 7002 Δ*addAB*. Next, both HIP1 methylated and unmethylated pCMK179 and pCMK181 (Figure 7A) were transformed into PCC 7002 Δ*addAB*. In this case, if HIP1 were indeed functioning as a canonical Chi site, deletion of *addAB* should negate any positive impacts of HIP1 inclusion/methylation on transformation efficiency. However, as seen in Figure 7B, even in PCC 7002 Δ*addAB*, transformation efficiencies were similarly improved (by up to ~130-fold, compared to ~128-fold in wild-type PCC 7002; Figure 3) when a homologous HIP1 site was included and methylated. That said, definitive conclusions cannot be drawn here since it is possible that additional or alternative nuclease/helicase complexes are instead associated with any putative Chi site function of HIP1 in PCC 7002. Nevertheless, *addAB* was still found to be a crucial component of natural transformation in PCC 7002 since, relative to wild-type, the total number of transformants obtained for both pCMK179 and pCMK181 were nearly 400- to 500-fold lower for PCC 7002 Δ*addAB* (Supplemental Figure 3).

**Figure 7.**
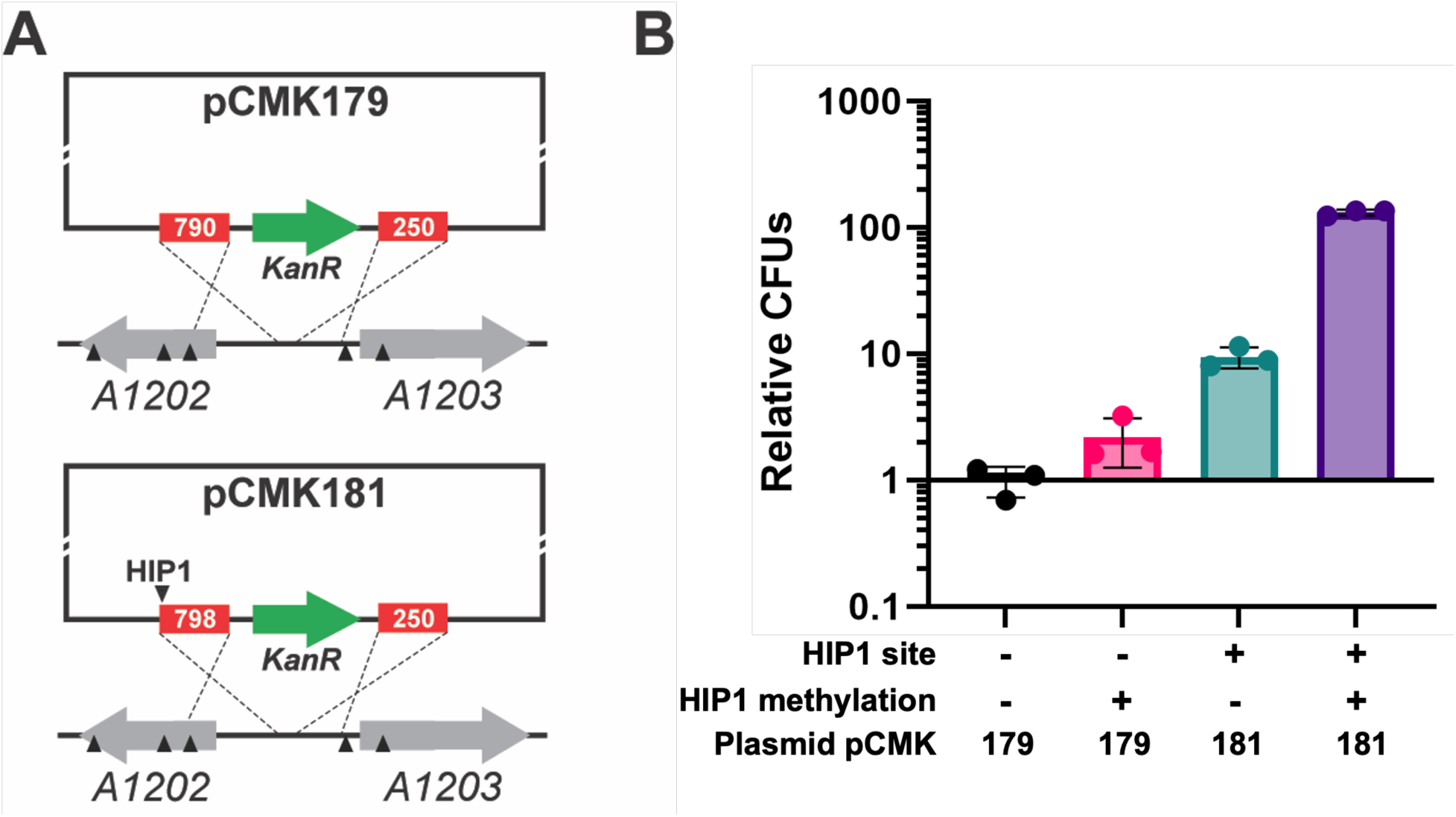
Investigating the potential function of HIP1 as a cyanobacterial Chi site analogue in PCC 7002 ΔaddAB. A: Schematic illustrating the number and positioning of homologous HIP1 sites (if any) located in the homology arms of pCMK179 and pCMK181. Both plasmids integrate a Kan^R^ cassette into the NS2 chromosomal locus. Solid black triangles represent native and homologous HIP1 sites. B: Effects of including and methylating a homologous HIP1 site on the relative number of colony forming units (CFUs) obtained. For both plasmids, CFUs were first normalized by dividing by the volume of cells plated and mass of DNA transformed. ‘Relative CFUs’ were determined by comparing the normalized results to those obtained for pCMK179. Error bars represent standard deviation from triplicate experiments.

## Discussion

Here we demonstrate how natural transformation efficiencies in PCC 7002 are enhanced by including a HIP1 site within the homology arms of a chromosomal integration plasmid. Moreover, when the homologous HIP1 site is also methylated by HIP1 methyltransferase, the overall enhancement can approach at least ~100-fold. Meanwhile, natural transformation efficiencies were also improved (albeit to a lesser extent) when a non-homologous HIP1 site is placed elsewhere on the plasmid backbone, but only when that site is also methylated. This was found to be true in the case of both chromosomally integrating and episomal replicating plasmids. These differing outcomes could be taken to suggest that HIP1 serves not one but multiple, distinct roles during natural transformation and/or integration. While the practical outcomes towards improving transformation efficiency are clear, the precise function of HIP1 in PCC 7002 currently remains elusive. However, as evidenced by Figure 2, since the addition of an unmethylated HIP1 site did not reduce transformation efficiency but instead improved it, it can be concluded that: *i)* HIP1 is not part of a native restriction enzyme system in PCC 7002 and, as such, *ii)* the improvements demonstrated here are not simply due to blocking such a system (as previously suggested in PCC 6803^15^). On the contrary, unlike known restriction sites, the consensus HIP1 site and related variants instead show a positive selection bias^12,13^. Furthermore, in PCC 7002, native HIP1 sites are periodically rather than randomly distributed throughout the chromosome; a feature that instead hints at its potential role in the maintenance and/or repair of chromosome structure^12^. Since cyanobacteria are constantly exposed to DNA damaging UV radiation and photosystems-generated reactive oxygen species (ROS)^32^, mechanisms for improving or stimulating DNA repair carry clear advantages.

Unlike when unmethylated, methylated HIP1 sites increased transformation efficiency when located in both homologous and non-homologous regions of both integrating and replicating plasmids. This position-independent behavior suggests that methylated HIP1 site recognition may occur prior to and separately from recombination, potentially during DNA binding or uptake; as would be the case for a DNA uptake sequence. DNA uptake sequences are common across naturally transformable, pathogenic bacteria, where they promote increased transformation of incoming DNA containing the sequence, allowing cells to discriminate between foreign DNA and that of the host or related bacteria^33,34^. Interestingly, while most DNA uptake sequences are non-palindromic (unlike HIP1), *Campylobacter jejuni* utilizes a palindromic DNA uptake sequence (RAATTY), thus leaving the door open to this prospect in PCC 7002. In *C. jejuni*, the RAATTY sequence is repeated throughout its genome (occurring every 60 bp on average) and it also contains an orphan methyltransferase that targets it^35^. Moreover, it has been shown that only a methylated RAATTY sequence serves as a DNA uptake signal in *C. jejuni*; with DNA containing a methylated RAATTY sequence displaying higher transformation efficiencies than if unmethylated or lacking the sequence altogether^35^. Lastly, beyond initial uptake, DNA uptake sequences have also been reported to positively impact natural transformation via as yet unidentified, secondary mechanisms (i.e., post-uptake)^36^.

Finally, the contrasting outcomes observed between Figures 2 and 3 vs. 5 and 6 indicate that, although unmethylated HIP1 sites can also increase transformation efficiency, this occurs only when they are included in regions sharing homology with the chromosome; an outcome that could alternatively suggest that HIP1 functions as a recombination hotspot or Chi site. However, since deletion of *addAB* did not abrogate the transformation efficiency increase gained by including a homologous HIP1 site (Figure 7), this suggests that HIP1 is at least unlikely to be a canonical Chi site (note: known Chi sites are also not palindromic or methylated). Of course, this does not exclude HIP1 from functioning as an alternative type of recombination hotspot that proceeds by another, distinct mechanism. Indeed, it is possible that a different, as yet unidentified nuclease/helicase complex (e.g., RecFOR homolgue^37,38^ or MutS/MutL/MutH/UvrD complex^39^) may be responsible for mediating such a potential Chi-like effect in PCC 7002.

Taken together, we postulate that HIP1 improves transformation through two separate mechanisms, the combination of which enables transformation efficiency to be increased by >100-fold. In the case of homologous HIP1 sites, both methylated and unmethylated, improve transformation efficiency by acting as Chi-like sites; promoting and perhaps even directly aligning homology-based DNA repair. In addition, our collective results further show how methylated HIP1 sites act in a position independent manner for both integrating and replicating plasmids and thereby may serve as a mechanism for cyanobacteria to discriminate between foreign versus cyanobacterial DNA during natural transformation (e.g., during DNA binding, uptake, or otherwise upstream of recombination). To truly discern the function of HIP1, however, additional mechanistic studies are needed. First, a comprehensive knockout library of putative PCC 7002 recombination proteins would aid in more rigorously investigating the potential function of HIP1 as a putative Chi site. Meanwhile, DNA affinity purification^40^ could be used to identify any proteins that specifically bind to HIP1 sites. The corresponding genes could then be knocked out to determine if they are indeed responsible for the observed increases in HIP1-mediated transformation efficiency. Finally, labeled DNA uptake assays could be performed on plasmids with and without HIP1 sites, since known DNA uptake sequences have been shown to increase rates of uptake rate^35,36^.

## Conclusion

Here we present an easily implemented approach for improving transformation efficiencies of chromosomal integration plasmids by nearly 100-fold in PCC 7002 by including and methylating a single homologous HIP1 site. For the greatest effect, at least two HIP1 sites should be located within both homology arms and methylated by HIP1 methyltransferase using *E. coli* DHIP1 or similar strain. Meanwhile, transformation of replicating plasmids can be improved nearly 10-fold by similarly including a methylated HIP1 site. These strategies show promise for applications requiring high transformation efficiencies, such as library construction and screening. Considering the widespread distribution of the HIP1 sequence, we predict that the presented methods can also be applied to other, diverse cyanobacterial species. Our collective findings demonstrate that HIP1 is not part of a native restriction enzyme system in PCC 7002, but that this site instead enhances transformation efficiency via other yet unidentified mechanism(s), such as by stimulating of DNA recombination and/or altering of DNA uptake. Future studies are needed to more clearly understand the precise role and function of HIP1 in PCC 7002.

## Acknowledgements

This work was supported in part by a grant from the National Science Foundation (CBET-1705409). CK received financial support in the form of a Completion Fellowship from Arizona State University. We thank Tanya Falbel, Jonathan Lombardino, Prof. Briana Burton and Joshua Abraham from the University of Wisconsin-Madison for their helpful discussion and suggestions regarding the potential mechanism(s)/function of HIP1. We thank Christopher M. Jones for constructing the PCC 7002 Δ*addAB* strain, assistance with figures, and helpful discussions.

