## Supplemental Figures and Tables for "Highly Iterated Palindrome 1 (HIP1) sequence improves *Synechococcus* sp. PCC 7002 transformation efficiencies in a homology- and methylation-dependent manner"

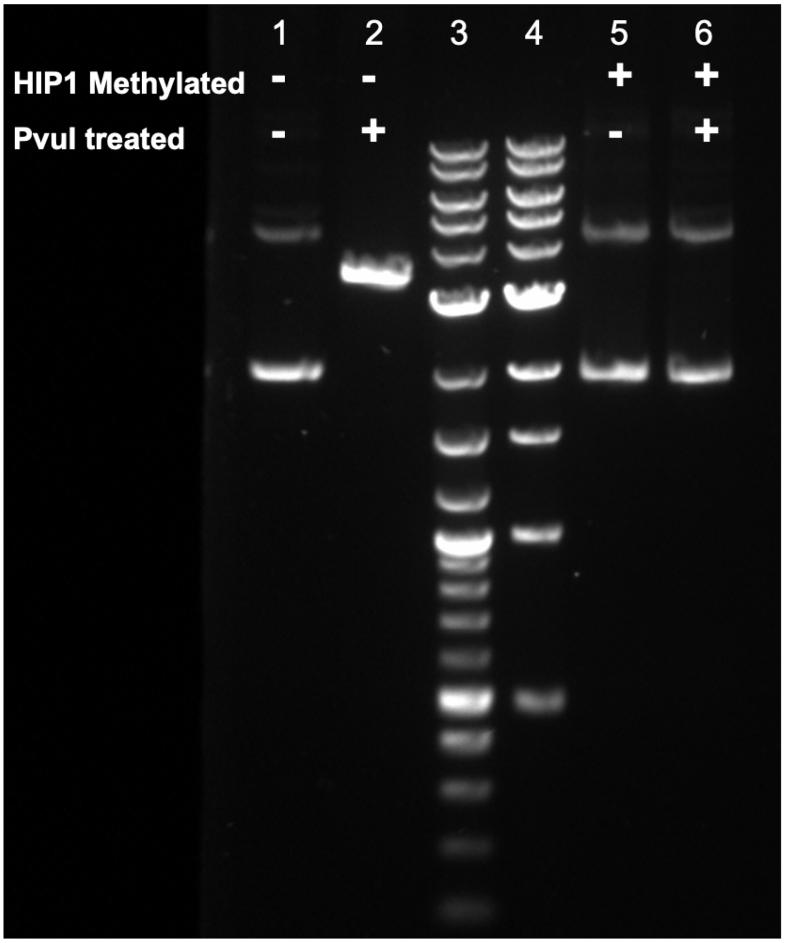


**Supplemental Figure 1.** *Assaying HIP1 methyltransferase activity in E. coli DHIP1. Methylation of the included HIP1 site on pCMK85 by HIP1 methyltransferase protects it from digestion by PvuI. pCMK85 only contains one HIP1 site and is 3382 bp Lane 1: pCMK85 miniprepped from DH10β (dam+ HIP1-) without PvuI. Lane 2: pCMK85 miniprepped from DH10β (dam+ HIP1-) with PvuI. Lane 3: 1 Kb Plus DNA ladder (NEB). Band sizes (in kb; top to bottom): 10, 8, 6, 5, 4, 3, 2, 1.5, 1.2, 1.0, 0.9, 0.8, 0.7, 0.6, 0.5, 0.4, 0.3, 0.2, 0.1. Lane 4: 1 Kb DNA ladder (NEB). Band sizes (in kb; top to bottom): 10, 8, 6, 5, 4, 3, 2, 1.5, 1.0, 0.5. Lane 5: pCMK85 miniprepped from DHIP1 (dam+ HIP1+) without PvuI. Lane 6: pCMK85 miniprepped from DHIP1 (dam+ HIP1+) with PvuI.*

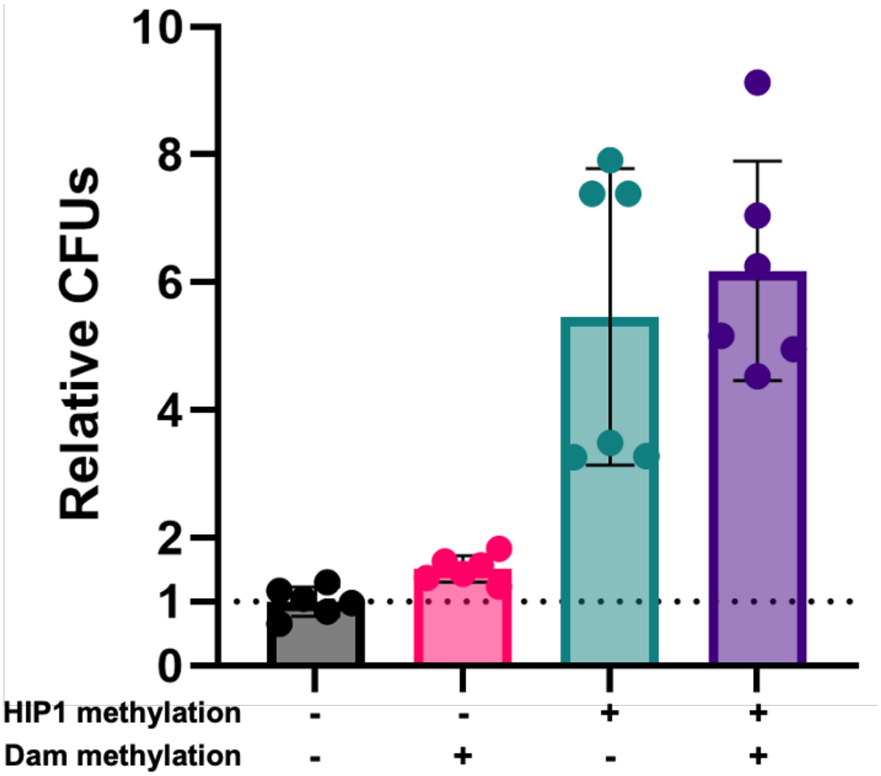


**Supplemental Figure 2.** *Comparing transformation efficiencies when delivering Dam and/or HIP1 methylated plasmids to PCC 7002. Plasmid pCMK85 with both unmethylated HIP1 and Dam sites were miniprepped from dam-/dcm- E. coli (NEB). To obtain plasmids with only methylated Dam sites (and not HIP1), plasmids miniprepped from dam-/dcm- E. coli were treated in vitro with Dam Methyltransferase (NEB). To obtain plasmids with only methylated HIP1 sites (and not Dam), the plasmid of interest was co-transformed with pAC-Cv into dam-/dcm- E. coli, then isolated from the miniprepped mixture via gel extraction. Plasmids with both methylated HIP1 and Dam sites were alternatively generated via in vitro treatment of the above gel extracted plasmids with Dam Methyltransferase. CFUs were first normalized by dividing by the volume of cells plated and mass of DNA transformed. 'Relative CFUs' were determined by comparing the normalized results to those obtained for pCMK85 miniprepped from dam-/dcm- E. coli. Error bars represent standard deviation from six experiments. Note that the total transformants obtained*

*for dam+ HIP1- pCMK85 was only 1.5 ± 0.1-fold greater than that of dam- HIP1- pCMK85, whereas no significant difference was seen between dam+ HIP1+ pCMK85 and dam- HIP1+ pCMK85 (6.1 ± 1.5 and 5.4 ± 2.1 Relative CFUs, respectively).*

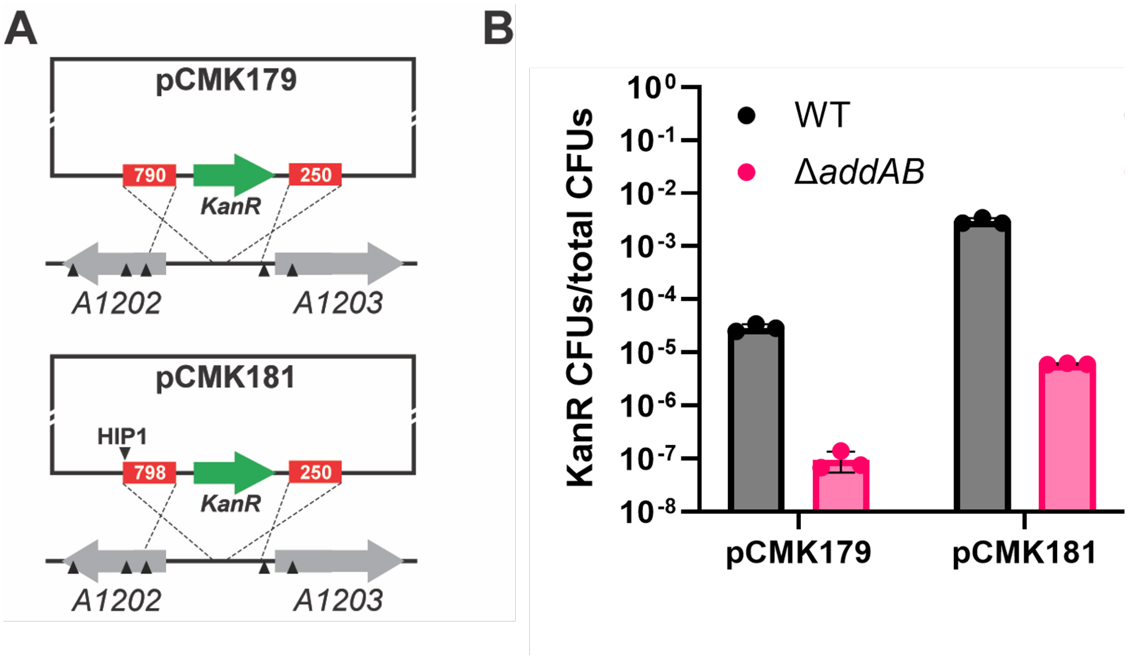


**Supplemental Figure 3.** *Effect of addAB deletion on transformation efficiency in PCC 7002. A: Schematic illustrating the number and location of homologous HIP1 sites (if any) in pCMK179 and pCMK181. Both plasmids integrate a $Kan^R$ cassette into the NS2 chromosomal locus. B: Effect of deleting addAB on transformation of pCMK179 and pCMK181 in PCC 7002. Data shown are compiled from Figure 4 (transformation into PCC 7002) and Figure 5 (transformation into PCC 7002 ΔaddAB). 'KanR CFUs/total CFUs' measured as the number of Kanamycin-resistant colonies divided by the total number of colony forming units (CFUs). Error bars represent standard deviation from triplicate experiments.*

**Supplemental Table 1.** *Primers designed and used in this study.*

| Primer name | Sequence | Used to construct |
|---|---|---|
| ck385 F | CTACAGAACATCAGACCTTGAAGACATGTTTTAGAGCTAGAAATAGCAAG TTAAAATAAG | pKDsgRNA gidB-3 |
| ck386 R | TTCAAGGTCTGATGTTCTGTAGAGTTGAAGACTAGTGCTCAGTATCTCTAT CACTGATAG | pKDsgRNA gidB-3 |
| ck463 F | TGACCTCACCTGCTTGAATGGCTAACTGCTTTAGGAATGGGATTTG | pCMK83 |
| ck464 R | TGACCTCACCTGCGGATGTATGTCAGCCCCATACGATATAAGTTG | pCMK83 |
| ck533 F | CAAGACCACCTGCTATTCCGTGAGTTTTCGTTCCACTGA | pCMK84 |
| ck534 R | CTGTATCACCTGCTACTTGCCCTCGGTTTCCTGGACTAAC | pCMK84 |
| ck535 F | CTAGACCACCTGCAACTGGCACCTCGCTAACGGATTCA | pCMK84 |
| ck536 R | CTAGACCACCTGCTGAGACGGGGCGTAATTTTTTTAAGGCAGT | pCMK84 |
| ck537 F | TATTGACACCTGCTAGACCGTGAGTTTTCGTTCCACTGA | pCMK85 |
| ck538 R | TAACATCACCTGCTAATATCGCCCTCGGTTTCCTGGACTAAC | pCMK85 |
| ck539 F | ATATTCCACCTGCAATACGATCGCCACCTCGCTAACGGATTCAC | pCMK85 |
| ck540 R | TAGACCACCTGCTCTAACGGGGCGTAATTTTTTTAAGGCAGT | pCMK85 |
| ck541F | GTCAATCACCTGCACAATCGCCAGACCCCGTAGAAAAGATCAAAG | pCMK95 |
| ck542 R | GTCAATCACCTGCATATGCGATCGCTCAGTGGAACGAAAACTCA | pCMK95 |
| ck590 R | GTTCTCCACCTGCACTACGATCGCTCACTCAAAGGCGGTAAT | pCMK100 |
| ck589 F | GTTCTACACCTGCACTTATCGCGATTCATTAAAATCTCAGGGATGG | pCMK100 |
| ck661 F | GAATACCACCTGCACCAGCTTCAAGCCCAAGGGTTTCGTAG | pCMK103 |
| ck662 R | GAATACCACCTGCTCCAGTCTGCCAGATCGAAAGAAAGAGGATC | pCMK103 |
| ck663 F | GAATACCACCTGCTCGTCATTGCTTCTTTTAACCTCAGATTAATTAGGC | pCMK103 |
| ck664 R | GAATACCACCTGCTCGTGCGATTGCGAATGTGATTGTAGAAGAATTG | pCMK103 |
| ck665 F | GAATACCACCTGCATTAGCTTATTGGGGGCTTGTTGAAGAT | pCMK104 |
| ck666 R | GAATACCACCTGCATTAGTCTAAGTCTTAATTTATGGGCATCTCC | pCMK104 |
| ck667 F | GAATACCACCTGCTTATCATTCCTTGTGATGATCACTACCGT | pCMK104 |
| ck668 R | TTAGACCACCTGCTAAAGCGAGTTTGAGGTAATTTTCGTTAATTCTAACC | pCMK104 |
| ck669 F | GAATACCACCTGCGGATGCTTGATTGGCTATGATCTACCAAAGCTG | pCMK105 |
| ck670 R | GAATACCACCTGCGGAAGTCTGGGTTAAATTAGGTCAAAAAAGTGTG | pCMK105 |
| ck671 F | GAATACCACCTGCACCACATTCCCCCTCTTGCCTACAGCAT | pCMK105 |
| ck672 R | GAATACCACCTGCACCAGCGAGAAACGAGATTATCTAAAACAGAAGCA | pCMK105 |
| ck817 F | ATCTTAGGTCTCCAGCCCAGAGTCCCGCTCAGAAG | pCMK138 |
| ck818 R | ATCTTTGGTCTCAGGCTTGGACTCCTGTTGATAGATCC | pCMK138 |
| ck945 F | ATCATTCACCTGCAAAGAAGGCAGCGTTGGGTCCTGGC | pCMK174 |
| ck946 R | ATCATACACCTGCTGAGCTGCTGGAGATGGCGGACGCG | pCMK174 |
| ck947 F | GCAGCCGCACGCGGCGCATCT | pCMK174 |
| ck948 R | CCTTAGATGCGCCGCGTGCGG | pCMK174 |
| ck949 F | ATCATTCACCTGCTATCTCGGGCACCTCGCTAACGGATTCA | pCMK175 |
| ck950 R | ATCATTCACCTGCTATCCCGAGCCTCGGTTTCCTGGACTAACAAG | pCMK175 |
| ck951 F | TTAGTTCACCTGCAATATCGGGCCTTGTGATGATCACTACCGTTTAAAAG | pCMK176 |
| ck952 R | ATCATTCACCTGCCCTTCCGAGTGCTTGGATTCTCACCAATAAAAAAC | pCMK176 |

| | | |
|---|---|---|
| ck961 F | ATCATTCACCTGCTTTACTTGACGTCGGAATTGCCAGCTG | pCMK179, pCMK180 |
| ck962 R | CAAATCCACCTGCACGATGAGTAATACGGTTATCCACAGAATCAGGG | pCMK179, pCMK180 |
| ck963 F | TACTTTCACCTGCCAGTCTCACCACAAGCCGGAGATTTG | pCMK179, pCMK180 |
| ck964 R | ATCATTCACCTGCCAGTCAAGTCTTAATTTATGGGCATCTCC | pCMK179, pCMK180 |
| ck965 F | AATCATCACCTGCACACCGATCGCCACCACAAGCCGGAGATTTG | pCMK181 |
| ck966 R | AACATTCACCTGCCTATATCGCCAGTAATACGGTTATCCACAGAATCAG | pCMK181 |
| ck965 F | AATCATCACCTGCACACCGATCGCCACCACAAGCCGGAGATTTG | pCMK182 |
| ck966 R | AACATTCACCTGCCTATATCGCCAGTAATACGGTTATCCACAGAATCAG | pCMK182 |
| CJ1564 | GACCGAGCGCGATCAAAAGCGAAAATGGTTTC | pCJ266 |
| CJ1565 | TGCAGATAACTTCGTATAGCATACATTATACGAACGGTAAAATTAAGTTACCCTCATCAC | pCJ266 |
| CJ1566 | CTTAATTTTACCGTTCGTATAATGTATGCTATACGAAGTTATCTGCAGCGGCCGCTACTA | pCJ266 |
| CJ1567 | CCAGTACCGTTCGTATAGCATACATTATACGAAGTTATACTAGAGCCAGGCATCAAATAA | pCJ266 |
| CJ1568 | TAGTATAACTTCGTATAATGTATGCTATACGAACGGTACTGGATTTGTGATGCCGGGAAG | pCJ266 |
| CJ1569 | AAATAGGCGTATCCCAGCATCCCCGATGATTTTC | pCJ266 |
| CJ1570 | CGGGGATGCTGGGATACGCCTATTTTTATAGGTTAATGTCATGATAATAATGGTTTCTTA | pCJ266 |
| CJ1571 | TTTTCGCTTTTGATCGCGCTCGGTCGTTCGG | pCJ266 |
